# FAK activity exacerbates disturbed flow-mediated atherosclerosis via VEGFR2-Cbl-NF-κB signaling

**DOI:** 10.1101/2024.12.06.627217

**Authors:** James M. Murphy, Duyen Thi Kieu Tran, Kyuho Jeong, Ly Nguyen, Mai Thi Nguyen, Dhananjay Tambe, Hanjoong Jo, Eun-Young Erin Ahn, Ssang-Taek Steve Lim

## Abstract

Atherosclerosis develops at predictable sites in the vasculature where branch points and curvatures create non-laminar disturbed flow. This disturbed flow causes vascular inflammation by increased endothelial cell (EC) barrier permeability and the expression of inflammatory genes such as vascular cell adhesion molecule-1 (VCAM-1). Vascular endothelial growth factor receptor 2 (VEGFR2) is important for flow-induced EC inflammation; however, there are still some gaps in the signaling pathway. Focal adhesion kinase (FAK) is a protein tyrosine kinase whose expression has been implicated in flow-mediated signaling in ECs. However, the link between FAK and VEGFR2 in flow-mediated inflammation signaling has remained unelucidated. Here we found that priming of VEGFR2 with VEGF was critical for flow-mediated activation of FAK and NF-kB. Mechanistically, FAK activation triggers tyrosine phosphorylation of Casitas B-lineage lymphoma (CBL; an E3 ubiquitin ligase) that interacts with VEGFR2 under flow conditions. Further, *Apoe-/-* mice fed a western diet (WD) exhibited increased FAK activity within the atheroprone disturbed flow region of the inner aortic arch compared to the outer arch. Disturbed flow-induced FAK activation is associated with elevated VEGFR2 on the surface of ECs of the inner aortic arch, but not in the outer arch. Taken together, these data suggest that suppression of augmented FAK activity under disturbed flow may prove beneficial in reducing pro-inflammatory signaling of the endothelial layer.

## Introduction

Atherosclerosis is a chronic inflammatory disease of the vessel wall, and atheroprone regions have been linked to abnormal hemodynamic factors (1). Branch points and curvatures of the vasculature are at a greater risk of developing atherosclerosis due to disturbed flow. Several studies have examined the gene expression profile in endothelial cells (ECs) exposed to either laminar or disturbed flow to better understand underlying causes in atherosclerosis development. These studies found that laminar flow decreases the expression of inflammatory and cell proliferation-associated genes, while disturbed flow increases the expression of these genes (2–5). Expression of inflammatory cell adhesion molecules (CAMs), such as vascular cell adhesion molecule-1 (VCAM-1), intercellular adhesion molecule-1 (ICAM-1), and E-selectin, have been detected in humans and animals at both atheroprone regions and in atherosclerotic lesions (6,7). Disturbed flow also increases the expression of monocyte chemoattractant protein-1 (MCP-1) and promotes monocyte attachment to ECs (8). To better understand the molecular mechanisms behind disturbed flow-induced inflammation and atherosclerosis, mouse models of disturbed flow have been developed. The gold standard in studying disturbed flow and atherosclerosis is the partial carotid artery ligation (PCL) in apolipoprotein E -/- (*Apoe-/-*) mice fed an atherogenic diet (9). The PCL model is beneficial in that atherosclerotic plaques develop within the carotid artery in 2-3 weeks, as opposed to the 12–24-week studies in normal atherosclerosis mouse models.

Nuclear factor-κB (NF-κB), a transcription factor closely linked with inflammatory molecule expression, is thought to play a key role in the progression of atherosclerosis. In ECs, expression of NF-κB and inhibitor of NF-κB α (IκBα) were increased in atheroprone regions compared to atheroprotective regions (10). Low-density lipoprotein (LDL) receptor -/- (*Ldlr-/-*) or *Apoe-/-* mice fed an atherogenic diet had increased nuclear NF-κB and phosphorylated IκBα in atheroprone regions (10,11). In human ECs, disturbed flow increases phosphorylation of NF-κB and NF-κB nuclear translocation, and laminar atheroprotective flow decreases nuclear NF-κB localization (12,13). Some of the upstream signaling events required to activate NF-κB in response to changes in flow have been elucidated. Shear stress induces expression of both vascular endothelial growth factor (VEGF) and VEGF receptor 2 (VEGFR2) (14–16). VEGFR2 acts as a mechanotransducer for flow-mediated NF-κB activation (17,18). Both nonselective tyrosine kinase inhibitors and VEGFR2 kinase inhibition blocked NF-κB activation in response to shear stress (16,18,19).

An important scaffolding protein in VEGFR2 flow-mediated signaling is Casitas B-lineage Lymphoma (CBL). CBL contains several tyrosine residues that are phosphorylated following flow (17) and contains a tyrosine kinase binding domain that allows it to bind to phosphotyrosine residues (20). Flow induces association of VEGFR2 with Cbl (17), and tyrosine phosphorylation of CBL is important for flow-mediated activation of inhibitor of NF-κB (IκB) kinase β (IKKβ) (18). It is currently not known what kinase phosphorylates Cbl following the initiation of flow.

Another important mechanotransducer in flow-mediated NF-κB activation is the αvβ3 integrin. Studies have shown that activation of αvβ3 integrins is required for subsequent activation of VEGFR2 (17,21). Using a blocking antibody against αvβ3 integrin prevented VEGFR2-CBL association and reduced tyrosine phosphorylation of both (17). This study also showed that VEGFR2 activity was not required for αvβ3 integrin activation in response to flow, suggesting that αvβ3 integrin is upstream of VEGFR2 in flow-mediated NF-κB signaling (17). It is not currently known how αvβ3 integrin leads to association and tyrosine phosphorylation of VEGFR2 and Cbl.

Focal adhesion kinase (FAK) is an integrin-associated protein tyrosine kinase (PTK) that plays a key role in cell migration and proliferation. Studies have shown that FAK expression is required for flow-mediated activation of NF-κB (13,21). In FAK-/- mouse ECs, initiation of flow reduced phosphorylation of NF-κB at serine 536 (pS536) but did not change NF-κB nuclear translocation (13). Small interfering RNA (siRNA) against FAK also reduced flow-mediated phosphorylation of NF-κB in bovine aortic ECs (13), suggesting that FAK expression in flow-induced NF-κB activation is not limited to mouse ECs. FAK-/- mouse ECs failed to induce expression of ICAM-1 following initiation of flow (13), further supporting the importance of FAK expression in flow-induced NF-κB activation

Since FAK plays a key role in integrin signaling, a few studies have investigated the link between FAK and integrins in flow-mediated signaling. Inhibition of αvβ3 using a peptide antagonist prevented flow-induced activation of both FAK and NF-κB (21). Knockdown of either αv or β3 using siRNA was also able to reduce flow-induced activation of FAK and NF-κB, and this was correlated with reduction in expression of VCAM-1 and ICAM-1 (21). This study also showed that treatment with the αvβ3 peptide antagonist S247 or αv EC specific knockout was enough to reduce plaque size, VCAM-1 expression, and macrophage staining in PCL model of disturbed flow (21). These studies suggest that FAK activity be important for NF-κB activation in response to flow, however the molecular pathways linking FAK to NF-κB activation remain unknown.

Here we attempted to decipher a potential flow-mediated signaling pathway where FAK promotes crosstalk between CBL, and VEGFR2 to activate IKK and NF-κB. We show that FAK activity is important for phosphorylation of NF-κB and IKK following the initiation of flow. Importantly flow-mediated activation of both FAK and NF-κB requires the presence of VEGF, suggesting that activation of VEGFR2 signaling may be a prerequisite of flow-mediated FAK activation. In the PCL mouse model, we found FAK inhibition blocks lipid accumulation, VCAM-1 expression, and macrophage recruitment to the vessel walls.

## Results

### VEGF is required for flow-mediated activation of FAK and NF-κB

While VEGFR2 is important for flow-mediated NF-κB activation (17), it is not known whether priming of VEGFR2 with VEGF is required for flow-mediated activation of FAK and NF-κB. To test this, we performed a time course assay where human umbilical vein endothelial cells (HUVECs) were subjected to either disturbed flow (5 dynes/cm^2^) or laminar (12 dynes/cm^2^) with or without VEGF. In the absence of VEGF, both disturbed and laminar flow failed to increase phosphorylation of FAK (pY397) and NF-κB (pS536) (Fig. 1A and S. Fig. 1). However, cells in the presence of VEGF showed increased phosphorylation of both FAK and NF-κB (Fig. 1A and S. Fig. 1). Increased FAK and NF-κB activation was associated with corresponding increase in active IKKα/β (pS176/177) and loss of IκBα (Fig. 1A and S. Fig. 1). While FAK expression is important for NF-κB phosphorylation (13), it is not known if FAK catalytic activity is required for NF-κB activation. Pretreatment with a small molecule FAK inhibitor (FAK-I, PF-271) prevented FAK activation, and blocked NF-κB activation in response to disturbed flow (Fig. 1B). These data suggest that VEGF priming of VEGFR2 is required for flow-mediated signaling and that FAK activity is necessary to propagate the signal to the inflammatory NF-κB pathway.

**Figure 1.**
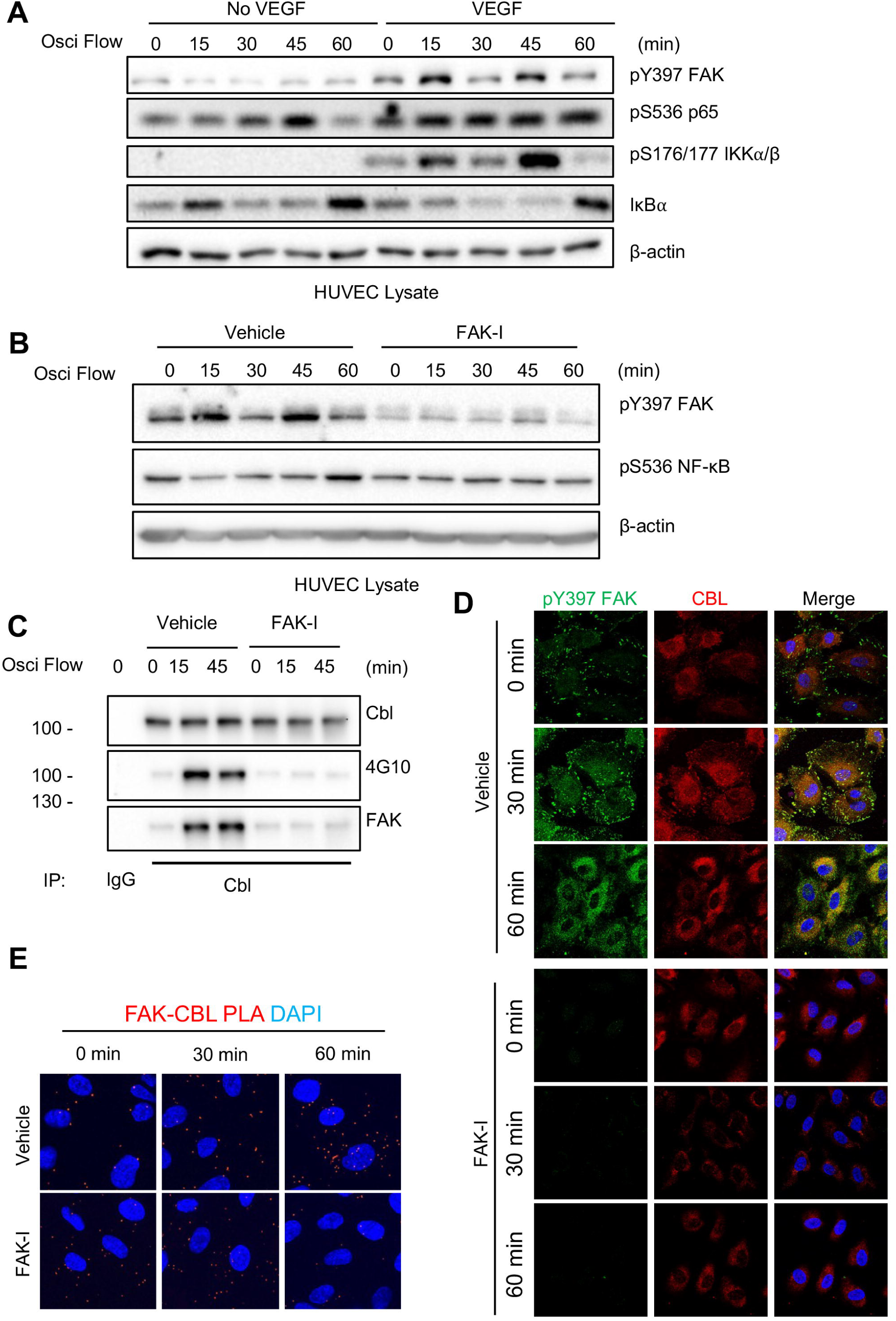
VEGF is required for flow-mediated formation of FAK-CBL complex and subsequent NF-κB activation in human endothelial cells. HUVECs were starved for 12 h in 0.2% FBS DMEM with or without VEGF (5 ng/ml) prior to initiation of disturbed flow (5 dynes/cm^2^). **(A)** Immunoblotting of active FAK (pY397), active NF-κB (pS536), active IKKα/β (pS176/177), IκBα, and β-actin as loading control are shown. **(B)** HUVECs were treated for 1 h with either DMSO or the FAK inhibitor (FAK-I, 2.5 μM) prior to initiation of disturbed flow (5 dynes/cm^2^). Immunoblotting shows active FAK (pY397), active NF-κB (pS536), and β-actin as loading control. **(C)** Immunoblotting of CBL immunoprecipitates (IP) for CBL, FAK, and Tyrosine phosphorylation (4G10). **(D)** HUVECs were treated with FAK inhibitor (FAK-I, 2.5 μM) prior to initiation of disturbed flow (5 dynes/cm^2^) for the indicated times. Immunostaining of pY397 FAK (Green) and CBL (Red). Nuclei were stained with DAPI (Blue). **(E)** HUVECs were treated with FAK inhibitor (FAK-I, 2.5 μM) prior to initiation of disturbed flow (5 dynes/cm^2^) for the indicated times. Proximity ligation assay (PLA) was performed using antibodies for FAK and CBL. Co- localization is indicated by red dots. Nuclei were stained with DAPI (Blue).

To better understand how FAK contributes to flow-mediated signaling via VEGFR2, we next examined CBL, a key E3 ubiquitin ligase and adapter protein known to be tyrosine phosphorylated and has been shown associated with VEGFR2 to mediate flow signaling in ECs (18,19,22). We performed CBL immunoprecipitation (IP) after 15- and 45-min disturbed flow, as these were the peak times in which we saw active FAK (Fig. 1A,B). disturbed flow increased CBL association with FAK and CBL tyrosine phosphorylation (Fig. 1C). However, pretreatment with FAK-I blocked flow-induced CBL-FAK interaction and tyrosine phosphorylation (Fig. 1C). To better understand CBL tyrosine phosphorylation and flow signaling, we overexpressed either wild-type (WT) or the triple tyrosine to phenylalanine mutant Y700/731/774F (Y3F) in HUVECs. Overexpression of WT CBL increased flow-mediated FAK activation (S. Fig. 2). However, overexpression of CBL Y3F blocked flow-mediated FAK activation (S. Fig. 2). These data suggest that while FAK activity is important for CBL tyrosine phosphorylation, CBL tyrosine phosphorylation also plays a role in maintaining FAK activity under disturbed flow. To better understand how disturbed flow might affect FAK and CBL, we performed immunostaining to observe their localization. Under static conditions active pY397 FAK stained primarily at the cell edge and focal adhesions (Fig. 1D). After 30 min of disturbed flow, pY397 FAK still showed staining on the cell edge but showed more cytoplasmic staining as well, which increased after 60 min (Fig. 1D). CBL showed primarily cytoplasmic localization under static conditions (0 min) and increased cell edge and focal adhesion localization at 30 min (Fig. 1D). Pretreatment with FAK-I reduced active pY397 FAK staining and prevented CBL relocalization following flow (Fig. 1D), suggesting that FAK activity is important for CBL recruitment to the cell edge following flow. We next performed proximity ligation assay (PLA) to evaluate co-localization of FAK and CBL. FAK-CBL showed increased interaction after 60 min flow in vehicle treated cells but was reduced by pretreatment with FAK-I (Fig. 1E). These findings suggest that disturbed flow increases FAK-CBL association, by which FAK phosphorylates CBL and CBL phosphorylation maintains FAK activity in ECs.

### FAK activity alters VEGFR2 cell surface expression

To better understand how FAK and CBL can affect flow-mediated signaling, we examined their roles on VEGFR2 expression and localization. Overexpression of the tyrosine phosphorylation deficient CBL Y3F mutant leads to increased VEGFR2 expression (S. Fig. 2), suggesting more stabilized expression of receptor which could be associated with reduced receptor endocytosis. Immunostaining of VEGFR2 also demonstrated that disturbed flow could alter VEGFR2 subcellular localization with elevated membrane expression observed after 60 min flow (Fig. 2A). FAK localization also changed to more cytoplasmic localization following 60 min flow (Fig. 2A). Pretreatment with FAK-I lead to increased FAK nuclear localization and prevented flow-mediated changes in VEGFR2 subcellular localization (Fig. 2A). Immunoblotting of CBL IP showed increased association with both FAK and VEGFR2 following disturbed flow (Fig. 2B), suggesting the three proteins form a complex. PLA also showed increased interaction between FAK-VEGFR2 and CBL-VEGFR2 following disturbed flow (Fig. 2C,D). However, pretreatment with FAK-I reduced FAK-VEGFR2 and CBL-VEGFR2 association after 60 min flow (Fig. 2C,D), suggesting that FAK activity is required for the formation of a VEGFR2-FAK-CBL signaling complex.

**Figure 2.**
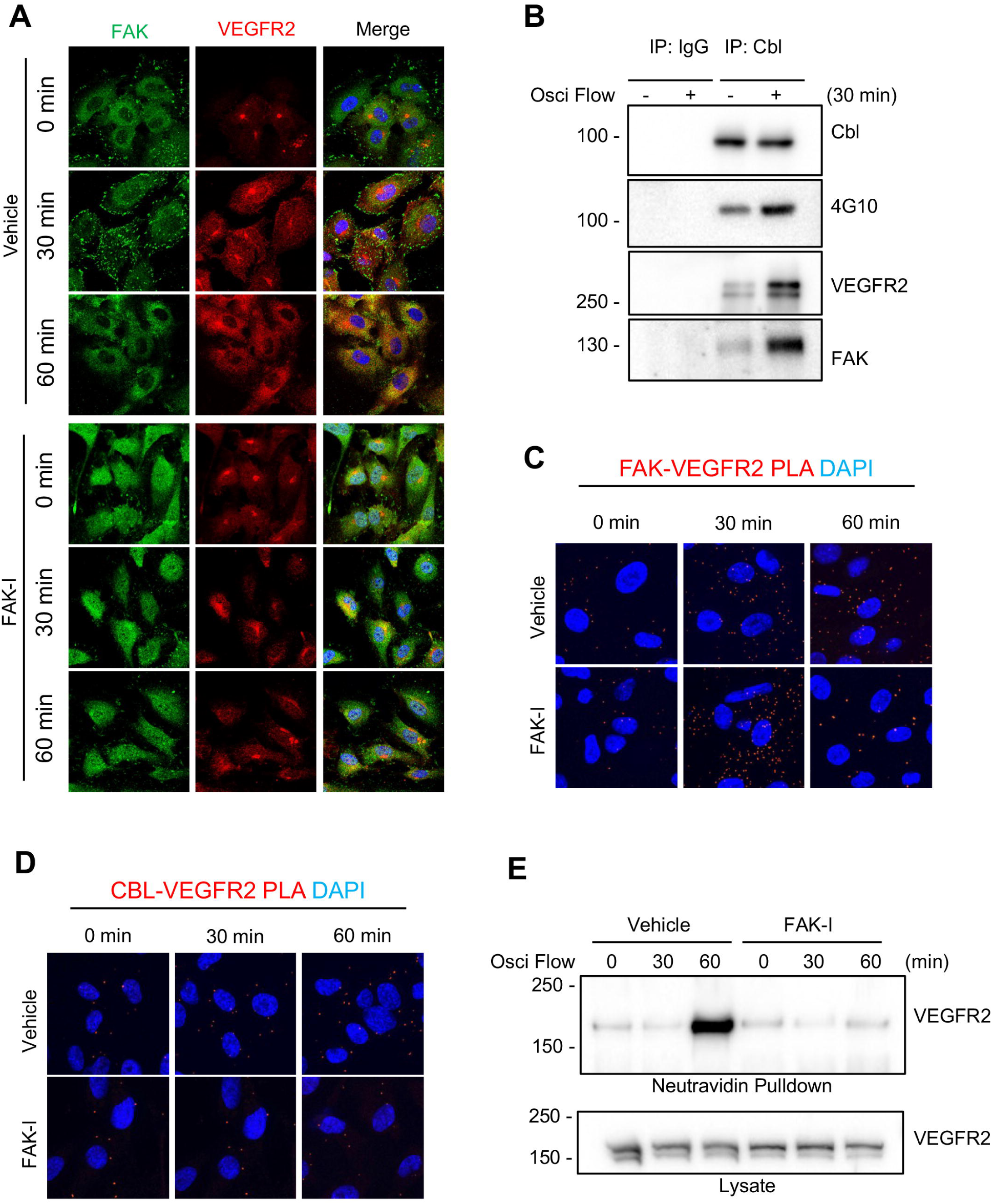
FAK activity is important for VEGFR2 expression on the plasma membrane. HUVECs were starved for 12 h in 0.2% FBS DMEM with or without VEGF (5 ng/ml) prior to initiation of disturbed flow (5 dynes/cm^2^). **(A)** HUVECs were treated for 1 h with either DMSO or FAK inhibitor (FAK-I, 2.5 μM) prior to initiation of disturbed flow (5 dynes/cm^2^). Immunostaining of FAK (Green) and VEGFR2 (Red). Nuclei were stained with DAPI (Blue). **(B)** Immunoblotting for CBL immunoprecipitates (IP) for CBL, VEGFR2, FAK and tyrosine phosphorylation (4G10). **(C)** HUVECs were treated with FAK inhibitor (FAK-I, 2.5 μM) prior to initiation of disturbed flow (5 dynes/cm^2^) for the indicated times. PLA was performed using antibodies for FAK and VEGFR2. Co-localization is indicated by red dots. Nuclei were stained with DAPI (Blue). **(D)** HUVECs were treated with FAK inhibitor (FAK-I, 2.5 μM) prior to initiation of disturbed flow (5 dynes/cm^2^) for the indicated times. PLA was performed using antibodies for CBL and VEGFR2. Co-localization is indicated by red dots. Nuclei were stained with DAPI (Blue). **(E)** HUVECs were treated for 1 h with either DMSO or the FAK inhibitor (FAK-I, 2.5 μM) prior to initiation of disturbed flow (5 dynes/cm^2^) for the indicated times. Immunoblotting of pulldown of biotin labeled surface proteins for VEGFR2.

Receptor endocytosis and sorting to either lysosomes or recycling endosomes play an important role in growth factor signaling (23). As Cbl has been shown to promote VEGFR2 internalization and FAK activity appears to be important for Cbl and VEGFR2 redistribution following flow, we examined VEGFR2 association with the endocytic pathway. VEGFR2 showed increased colocalization with early endosome antigen 1 (EEA1, an endosomal marker) (S. Fig. 3). However, pre-treatment with FAK-I blocked localization of VEGFR2 with EEA1 (S. Fig. 3), suggesting FAK activity is required for VEGFR2 localization to the endocytic compartments. To evaluate if FAK activity altered the surface expression of VEGFR2 under disturbed flow, proteins on the cell surface were biotinylated after being subjected to flow. After 60min of flow, there was a large increase in VEGFR2 surface expression which was blocked by FAK-I (Fig. 2E). These data suggest that FAK activity is important for the relocalization and recycling of VEGFR2 to the cell surface under disturbed flow.

### FAK inhibition reduces lipid accumulation and vascular inflammation in a mouse model of disturbed flow

We next looked at in vivo areas of disturbed or laminar flow to see if there were differences in FAK activity, Cbl, or Vegfr2. *Apoe-/-* mice underwent partial carotid ligation (PCL) surgery and were fed a Western diet (WD) for 2 weeks. Mice were treated with either vehicle or FAK-I twice daily for the duration of the experiment. First, we examined the outer and inner regions of the aortic arch, which are known to undergo laminar and disturbed flow, respectively. The inner aortic arch showed increased active pY397 FAK staining compared to the outer arch (Fig. 3A). Similarly, Cbl and Vegfr2 showed elevated staining within the inner arch compared to the outer (Fig. 3B,C). Mice treated with FAK-I showed reduced active pY397 FAK in both regions of the aortic arch (Fig. 3A). While there was no change in Cbl expression within the inner arch (Fig. 3B), Vegfr2 showed decreased expression in both the inner and outer regions (Fig. 3C). These data suggest that the inner arch is more susceptible to a Vegfr2-Cbl-FAK signaling axis than the outer arch.

**Figure 3.**
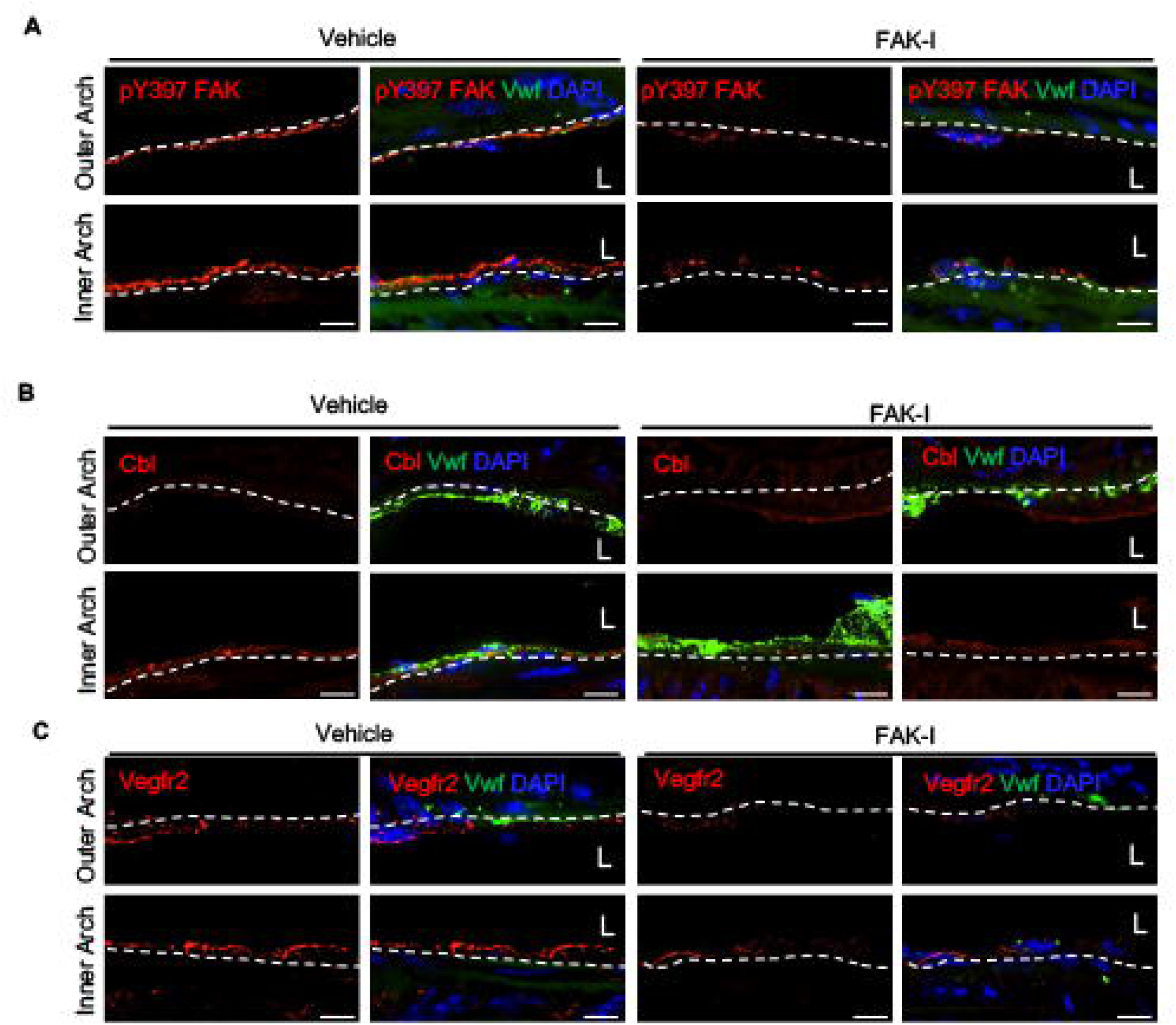
FAK shows increased cytoplasmic localization and activity at regions of disturbed flow. *Apoe-/-* mice underwent partial carotid ligation (PCL) and were treated with either vehicle or FAK inhibitor (FAK-I, 35 mg/kg) while on a western diet (WD) for 2 weeks. The outer and inner aortic arches were immunostained for **(A)** active pY397 FAK (Red), **(B)** Cbl (Red), or **(C)** Vegfr2 (Red). Endothelial cells were stained with vWF (Green) and nuclei with DAPI (Blue).

We next analyzed the left carotid artery that underwent PCL. Lipid staining with Oil Red O showed strong lipid accumulation in vehicle-treated arteries compared to FAK-I treated (Fig. 4A). No Oil Red O staining was observed in sham arteries (Fig. 4A). FAK-I efficacy was evaluated via immunoblotting of heart lysates for active pY397 FAK (S. Fig. 4A). Immunostaining of ligated arteries showed that FAK-I treatment reduced the high levels of active pY397 FAK that were observed in vehicle controls (Fig. 4B). Further, vehicle-treated mice showed high expression of Cbl within the EC layer, but this expression was reduced in FAK-I mice (Fig. 4C). Vegfr2 expression was also increased in vehicle-treated mice compared to FAK- I (Fig. 4D), suggesting that disturbed flow elevates Cbl and Vegfr2. Using PLA, we also observed FAK-Vegfr2 and FAK-Cbl co-localization within ECs of injured arteries of vehicle-treated mice but not FAK-I (Fig. 4E,F). FAK-I treatment was able to reduce expression of Vcam-1 (Fig. 4B), active pS536 NF-κB (S. Fig. 4C) and macrophage recruitment (S. Fig. 4D) when compared to vehicle-treated mice. Taken together, these data support the notion that FAK inhibitors could be used to treat vascular inflammatory diseases such as atherosclerosis.

**Figure 4.**
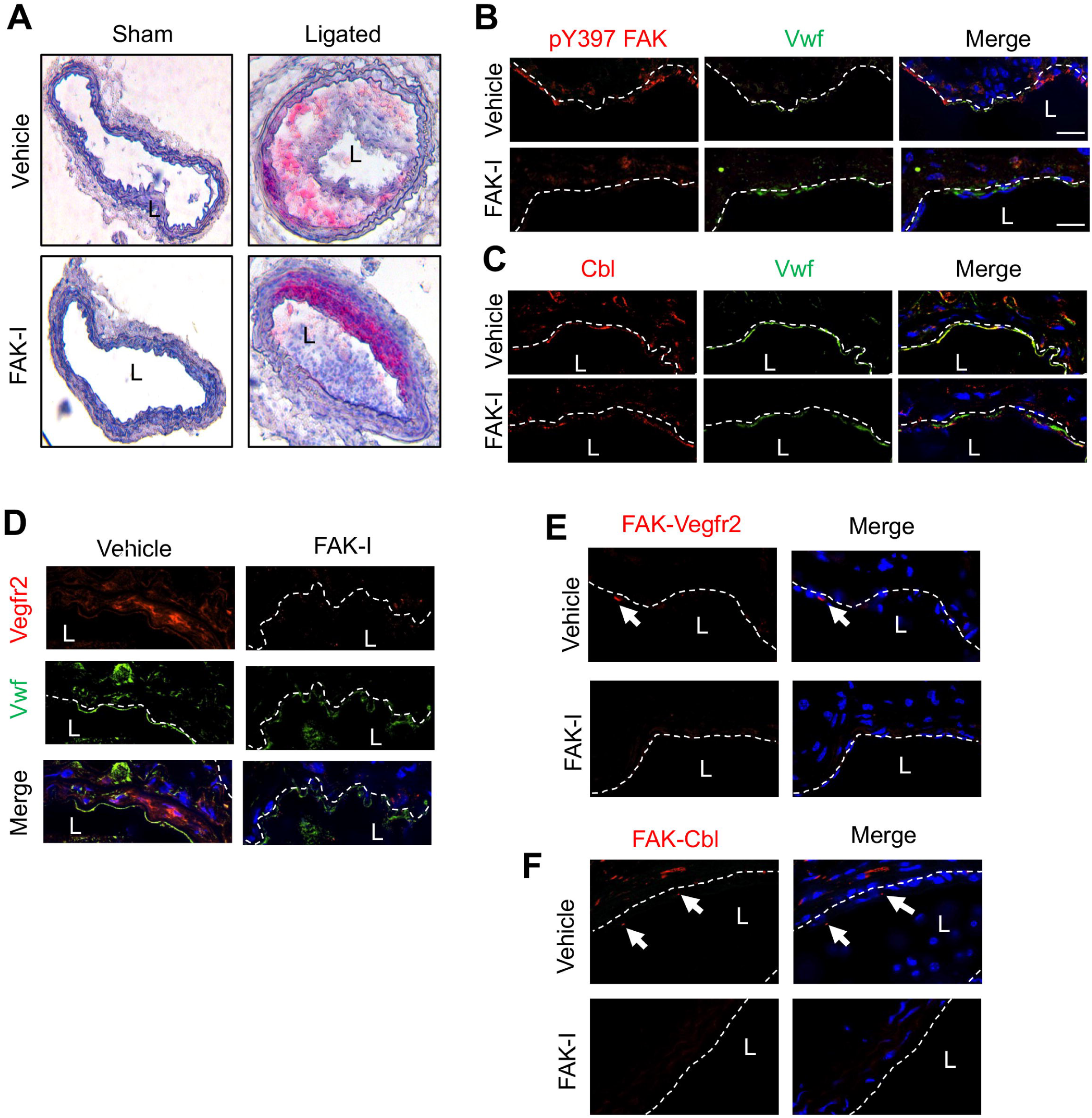
FAK inhibition reduces lipid accumulation in partial carotid ligation mouse model. Partial carotid ligation (PCL) was performed on *Apoe-/-* mice and were treated with vehicle or FAK inhibitor (FAK-I, 30 mg/kg, twice daily) while on a western diet (WD) for 2 weeks. **(A)** Representative images of Oil Red O staining of sham and ligated carotid arteries are shown. Immunostaining of carotid arteries for **(B)** pY397 FAK (Red), **(C)** Cbl (Red), or **(D)** Vegfr2 (Red). Endothelial cells were stained with Vwf (Green) and nuclei with DAPI (Blue). **(E and F)** PLA was performed with antibodies targeting **(E)** FAK and Vegfr2 or **(F)** FAK and Cbl. Red dots indicate positive co-localization. Nuclei stained with DAPI (blue).

### EC-specific FAK inhibition reduces PCL-induced remodeling and vascular inflammation

To determine if these changes were specific to FAK inhibition in ECs, we next used EC-specific FAK kinase dead (KD) genetic mouse model on the *Apoe-/-* background. FAK-WT and FAK-KD mice underwent PCL and were fed a WD for 2 weeks. Oil Red O staining of sham and ligated arteries revealed significant lipid accumulation and remodeling in FAK-WT mice, but not FAK-KD (Fig. 5A). FAK-KD mice showed reduced levels of pY397 FAK via immunostaining of ligated arteries for pY397 FAK (Fig. 5B). Interestingly, FAK-WT mice showed increased levels of CBL and VEGFR2 within ECs in ligated arteries (Fig. 5C,D) However, FAK-KD mice showed reduced Vegfr2 and Cbl immunostaining (Fig. 5C,D). FAK-WT mice also had elevated levels of Vcam-1 and active nuclear pS536 NF-κB in ECs compared to FAK-KD (S. Fig. 5A and Fig. 5D). Taken together, these data suggest that FAK activity is important to increase Vegfr2-Cbl expression and signaling to promote vascular inflammation under disturbed flow conditions.

**Figure 5.**
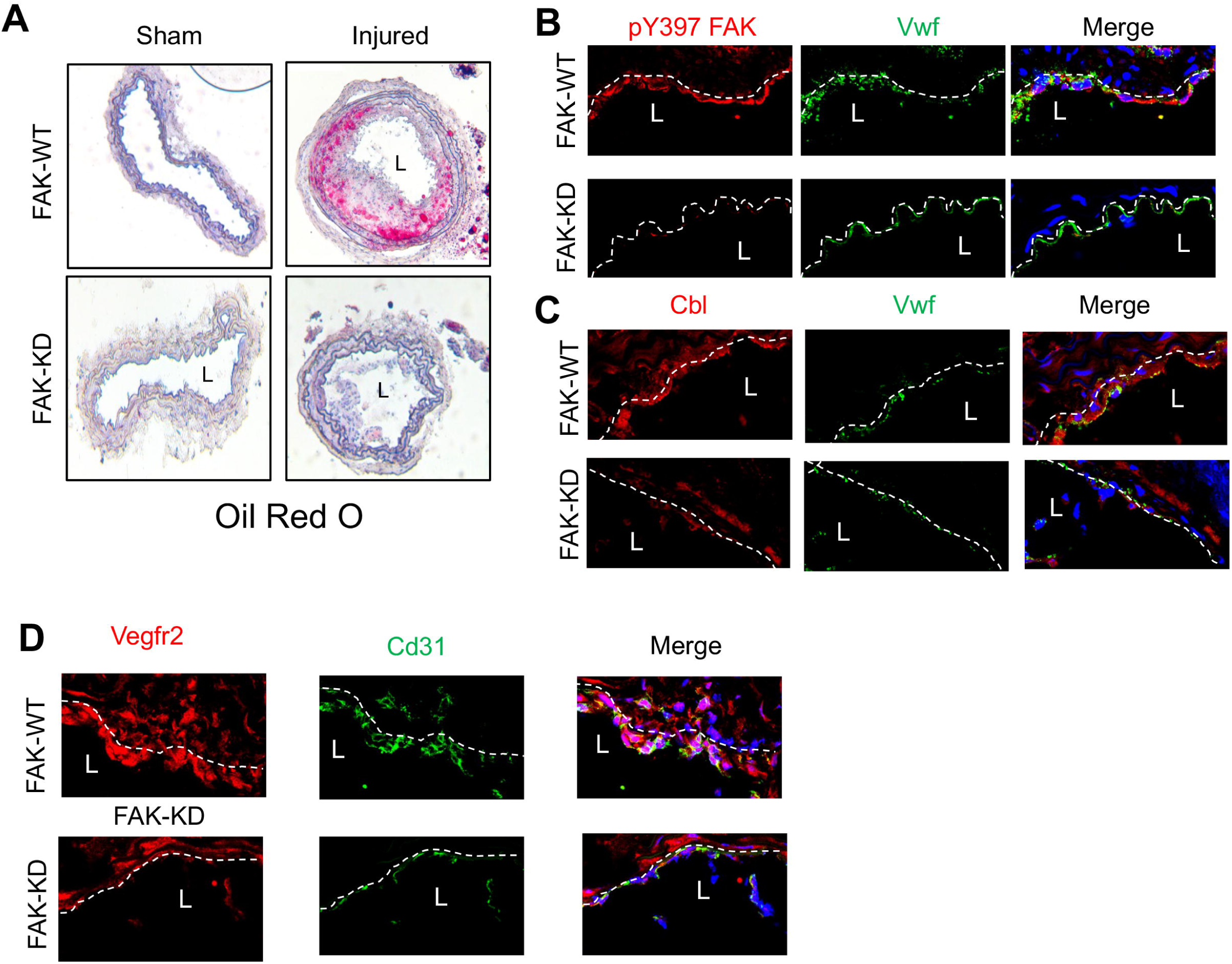
Endothelial cell specific FAK inhibition reduces disturbed flow-mediated plaque formation. Partial carotid ligation (PCL) was performed on *Apoe-/-;FAK-WT* and *Apoe-/-;FAK- KD* mice and fed a western diet (WD) for 2 weeks. **(A)** Representative images of Oil Red O staining of sham and ligated carotid arteries are shown. Immunostaining for **(B)** pY397 FAK (Red), **(C)** Cbl (Red), or **(D)** Vegfr2 (Red). Endothelial cells stained with Vwf (Green) and nuclei were stained with DAPI (blue).

**Figure 6.**
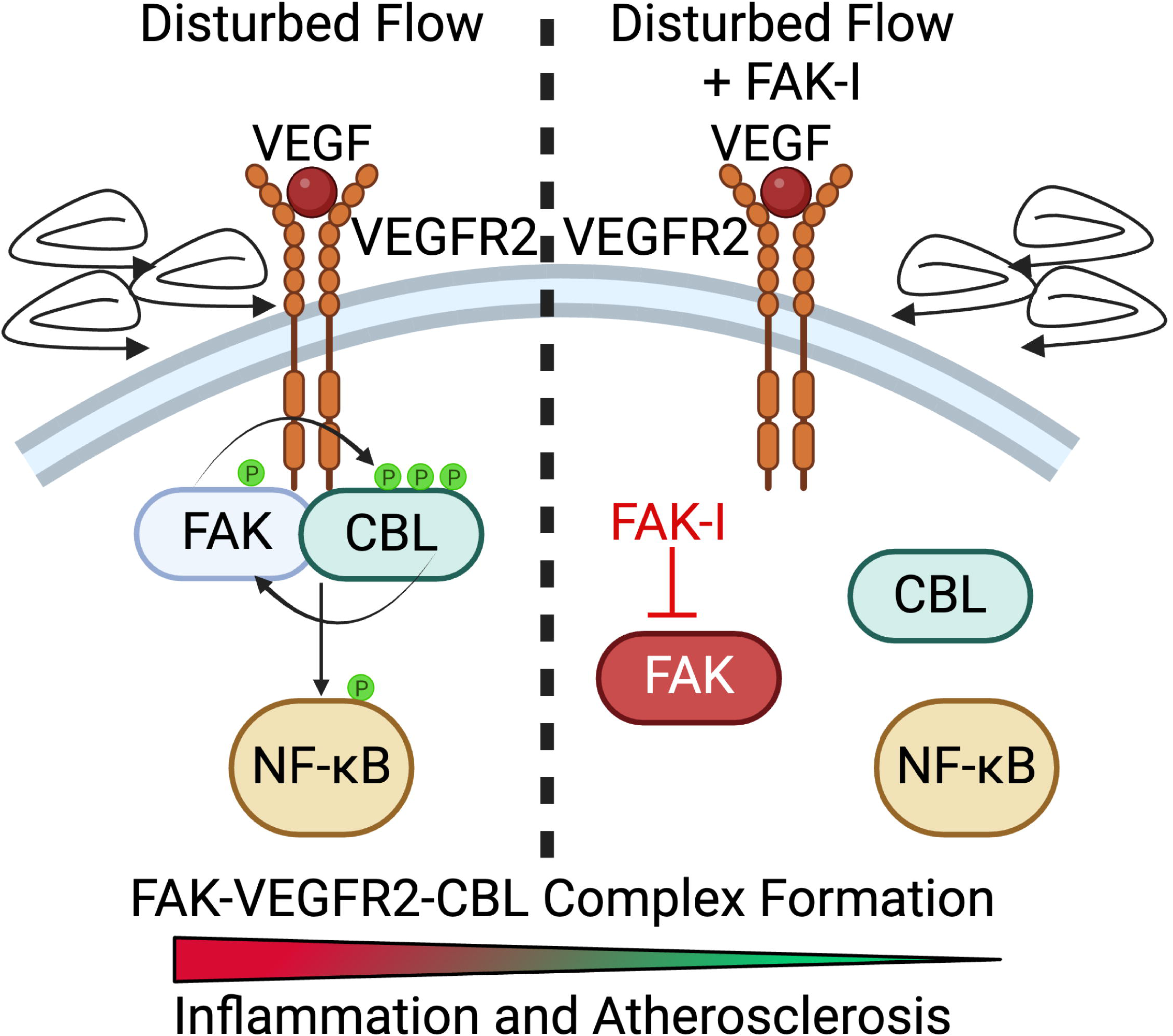
FAK works with Cbl-VEGFR2 complex to promote flow-mediated NF-kB activation and atherosclerosis progression. Proposed model of FAK translating integrin signaling to CBL-VEGFR2 complex under disturbed flow. The FAK-CBL-VEGFR2 complex then activates IKK-NF-kB pathway to promote inflammatory gene expression.

## Discussion

In the present study, we showed that FAK activity was important for flow-induced activation of NF-κB and IKK in ECs (Fig. 1B). Importantly, we found that FAK associates with CBL (Fig. 1C and E) and might co-localize with VEGFR2 (Fig. 2B,C). It has been shown that activation of αvβ3 integrin is required for both FAK activation (21) and VEGFR2 association with CBL (17). Those studies, along with our findings suggest that FAK activity and its localization may be important to connect integrin activation with VEGFR2- and CBL-mediated IKK activation.

VEGFR2 priming with VEGF was required for flow-induced activation of both FAK and NF-κB (Fig. 1A,B). CBL contains a SRC-holomolgy 2 (SH2) domain that allows it to bind to phosphotyrosine residues and act as a signaling scaffold (20), so it remains to be seen if prior phosphorylation of VEGFR2 is required for CBL to associate with VEGFR2 after initiation of flow. We showed that CBL IP pulled down CBL and was tyrosine phosphorylated, and the tyrosine phosphorylation was dependent on FAK activity (Fig. 1C). Interestingly, mutation of CBL tyrosine residues that are known to be phosphorylated following flow (18) was able to reduce FAK activation (S. Fig. 2). It appears that there might be a feedback loop between FAK activity and CBL tyrosine phosphorylation necessary to promote flow-mediated signaling into the cell.

VEGFR2 has been shown to undergo endocytic recycling under both stimulatory and nonstimulatory conditions (24,25). Inhibition of endosome-to-plasma membrane recycling resulted in increased VEGFR2 localization in a perinuclear compartment (25). We observed a similar perinuclear abundance of VEGFR2 in FAK-I treated cells (Fig. 2A). Rab4a was found to be important for VEGFR2 recycling and blocking VEGFR2 degradation upon VEGF stimulation (25). FAK inhibition was shown to reduce Rab5 and reduced early endosome formation in fibroblasts (26) and Rab5 was shown prevent VEGFR2 degradation (27). We saw a similar pattern in which FAK inhibition early endosome formation and blocked VEGFR2-EEA1 association (S. Fig. 3). Taken together, it appears that FAK activity is required for the endocytic-to-plasma membrane recycling and localization of VEGFR2 in ECs experiencing disturbed flow.

An important initiator in the development of atherosclerosis is disturbed flow within the vessel (1,9). While anti-hypertensives are used to lower blood pressure and reduce elevated shear stress along the vessel wall, they cannot change the native vascular structure which gives rise to disturbed flow at branch points and artery curvatures. Therefore, a therapy that can block the endothelial response to these native areas of disturbed flow could prove beneficial in slowing down atherosclerosis. Here we showed that small molecule pharmacological FAK inhibition was able to reduce lipid accumulation in a mouse model of disturbed flow (Fig. 4A) and reduced endothelial Vcam-1 expression (S. Fig. 4C) and macrophage recruitment (S. Fig. 4D). These data suggest that FAK inhibitors could be useful in blocking the endothelium response to disturbed flow.

Atheroprotective laminar flow promotes EC barrier stability and inhibits EC proliferation (28), which are important in preventing the accumulation of LDLs in the vessel wall. Atheroprone disturbed flow, however, promotes inflammatory gene expression, decreases EC barrier stability, and increases EC proliferation and turnover, making it more likely for LDLs to get embedded in the vessel wall (1–5). From EC-specific genetic kinase-dead FAK (KD-FAK) studies, it was shown that FAK activity was important for increasing EC barrier permeability (29). More work needs to be done to determine what role FAK has in ECs during either atheroprotective or atheroprone flow, and whether FAK inhibition can reduce disturbed flow-induced EC permeability.

## Materials and Methods

### Sex as a biological variable

Sex is not a variable when studying endothelial signaling changes to flow. As such, male and female mice data were combined.

### Animal Experiments

Animal experiments were approved by and performed in accordance with the guidelines of the University of South Alabama and University of Alabama at Birmingham Institutional Animal Care and Use Committees. Partial carotid artery ligation (PCL) was performed as explained elsewhere (9). Briefly, the left carotid artery (LCA) bifurcation is separated from the associated nerve and vein to reveal the four distal branches of the LCA: internal and external carotid arteries, occipital artery and superior thyroid artery. A 6-0 silk suture is used to ligate the internal carotid and occipital arteries in a single knot, while the external carotid artery will be ligated above the superior thyroid artery. In Sham controls, the same surgery was performed as in the injury model except for ligation of distal branches. The morning after PCL, FAK-I (PF-271, 30 mg/kg) or vehicle (30% [2-hydroxypropyl]-b-cyclodextrin/3% dextrose, Sigma) was administered by oral gavage twice daily till end of the experiment. Mice were also placed on high fat diet (Envigo, TD.88137) to accelerate atherosclerotic plaque formation.

### Collection of Tissue Samples

Mice were euthanized via IP injection of ketamine/xylazine. Mice were then perfused with 5 mM EDTA in PBS through the left ventricle. Next, mice were perfused with PBS, followed by 4 % PFA in PBS to fix tissue structure. The left and right carotids were isolated and fixed in optimal cutting temperature (O.C.T.) and frozen at −80 °C.

### Oil Red O Staining

Frozen blocks were allowed to reach −20 °C in cryostat prior to sectioning. Samples were cut at a thickness of 7 μm and were allowed to adhere to RT glass slides. Frozen sections were allowed to dry at RT for 10 min, and then fixed with 4 % PFA in PBS for 10 min at RT. Slides were then washed in PBS for 10 min. Tissues were incubated in propylene glycol for 10 min, then lipids were stained using 0.6 % Oil Red O in propylene glycol. Slides were washed three times for 15 min in 85 % propylene glycol at RT. Cover glass was attached and images were taken.

### Antibodies and Reagents

Anti-pY397 FAK (cat# 44-624G) was purchased from Life Technologies. Anti-FAK (clone 4.47), anti-phosphotyrosine (clone 4G10), and anti-GAPDH (clone 6C5) were purchased from Millipore. Anti-pS536 NF-κB (clone 93H1) and anti-pS176/188 IKKα/β (clone 16A6) were purchased from Cell Signal Technologies. Anti-β-actin (clone AC-74) was purchased from Sigma-Aldrich. Anti-VEGFR2 (clone C-1158), anti-IκBα (SC-371), anti-CBL (SC-1651), anti-vWF (SC-271409) were purchased from Santa Cruz Biotechnology. Anti-LAMP2 (ab5631) and anti-EEA1 (ab70521) were purchased from Abcam. Anti-FAK used for immunoprecipitation was homegrown. Recombinant human VEGF was purchased from Peprotech. The FAK inhibitor PF- 271 was purchased from MedKoo.

### Cell Culture

Human umbilical vein endothelial cells (HUVECs) were purchased from Lifeline Cell Technology and grown in VascuLife VEGF Endothelial Medium Kit (Lifeline Cell Technology).

### Flow Experiments

HUVECs were seeded onto fibronectin coated cover slips and allowed to reach confluency. Cells were then serum starved in 0.2 % DMEM for 12 h with or without VEGF (5 ng/ml) prior to flow. Experiments were performed using the FlexFlow and OsciFlow Flow Controller (FlexCell International) or orbital shaker. Cells were then subjected to either laminar flow (12 dynes/cm^2^) or disturbed flow (5 dynes/cm^2^) for indicated times. In this model, flow will be highest at the edge of the plate and decreasing the closer to the center. With laminar flow on the outside radius of the plate and disturbed shear stress experienced by the inner part of the plate. Shear stress was calculated as previously described (30):

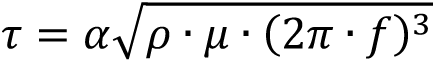

Where 1″ is the maximum shear stress (∼5 dynes/cm^2^), α is radius of orbit for the shaker (1.9 cm), ρ is the fluid density (1 g/cm^3^), μ is the fluid viscosity (8.94 x 10^-3^ g/(cm*sec), and *f* is the frequency of rotation (1.25 rounds/sec).

### Biotinylation Assay

Following flow, HUVECs were washed with cold PBS and then incubated with 2mM Sulfo-NHS-SS-11-Biotin (ApexBio, cat# A8015) for 30min on ice to label surface proteins. Cells were then washed with PBS containing 100 mM glycine three times. Cells were lysed in 1% Triton X-100 with protease inhibitor (Roche). Lysates were centrifuged for 1 h at 4°C, then incubated with neutravidin beads (Invitrogen) overnight with rotating at 4°C. Beads were then washed with 1% Triton X-100 three times. Biotinylated surface proteins were eluted by boiling in SDS loading buffer.

### Immunoblotting

Cells were lysed in RIPA buffer (pH 7.4) that included 4-(2-hydroxyethyl)-1- piperazineethanesulfonic acid (50 mM), sodium chloride (150 mM), Triton X-100 (1%), sodium deoxycholate (1%), sodium dodecyl sulfate (SDS;0.1%), glycerol (10%), and protease inhibitors (Complete Protease Inhibitor Cocktail, Roche, Mannheim, Germany). Lysates were cleared by centrifugation, and supernatants were boiled in SDS loading buffer. Samples were separated by SDS polyacrylamide gel electrophoresis (PAGE) and immunoblotted with indicated antibodies.

### Immunoprecipitation

Cells were lysed that includes Triton X-100 (1%), HEPES (50 mM), NaCl (150 mM), glycerol (10%), ethylene glycol tetra-acetic acid (1 mM), sodium pyrophosphate (10 mM), sodium fluoride (100 mM), sodium orthovanadate (1 mM), and protease inhibitors. Lysates were cleared by centrifugation, and equal amounts of proteins were subjected to immunoprecipitation with indicated antibodies. The lysates were rotated overnight at 4°C, then protein G or A agarose beads were added, and the mixture was rotated for 2 h at 4°C. The immunoprecipitates were washed three times with immunoprecipitation buffer and suspended with 2X SDS-loading buffer. Samples were separated by SDS-PAGE and immunoblotted with indicated antibodies.

### Immunocytochemistry

Cells were fixed using 4 % PFA in PBS for 10 min at RT and washed 3x with PBS for 5 min each. Cells were blocked and permeabilized in blocking buffer (10 % goat serum, 0.3 % Triton X-100, 1x PBS) for 45 min at RT. Slides were incubated overnight at 4°C with primary antibody in dilution buffer (1 % goat serum, 1 % BSA, 0.3 % Triton X-100, 0.01 % sodium azide, 1xPBS). Coverslips were washed 3 times in PBS for 15 min each at RT, and then incubated with Alexa Fluor secondary antibodies in the dark for 1 h at RT. Slides were washed 3 times in PBS for 15 min each at RT. Nuclei were stained using DAPI for 2 min, then washed in PBS for 5 min. Coverslips were mounted onto slides using Fluoromount-G (SouthernBiotech) and held in place using clear nail polish. Fluorescent microscopy was performed using a Nikon A1 Confocal microscope.

### Immunostaining

Frozen sections were fixed with cold acetone for 15 minutes and washed thrice with PBS. Samples were permeabilized with 0.1% Triton X-100 for 10 minutes, washed with PBS, and incubated with blocking solution containing 3% bovine serum albumin (BSA) and 1% goat serum for 1 hour at RT. Sections were incubated with primary antibodies overnight at 4°C. Sections were washed with PBS and incubated with conjugated goat anti-rabbit or mouse secondary antibodies (1:1000) (Alexa Fluor 594 or 488, Thermo Fisher) for 1 hour at RT. Slides were mounted (Fluoromount-G, SouthernBiotech, Birmingham, AL), and images were acquired with a confocal microscope at 60-fold magnification (Nikon A1R, Nikon, Tokyo, Japan).

### Proximity Ligation Assay

Cells or tissue sections were stained with the indicated antibodies and proximity ligation assay (PLA) was performed following the manufacturer’s instructions (DuoLink PLA, MilliporeSigma).

## Study Approval

Biosafety and IACUC

## Author Contributions

JMM, EYA, and STL conceived the idea. JMM, EYA, and STL designed the experiments. JMM, KJ, DTKT, LN, and MTN performed the experiments. DT and HJ provided training and resources. JMM, KJ, and STL analyzed the data. JMM and STL wrote the paper. JMM was assigned the first position in the authorship order since JMM initiated and contributed to the original idea of this work.

## Acknowledgments

We would like to thank Dr. Larry Samelson at the National Cancer Institute for kindly providing us with plasmids for Cbl-WT and Cbl-Y3F. This work was supported by American Heart Association 16GRNT30960007, National Institutes of Health R01HL136432 and HL158875 (to S. L.).

