## Supplemental Figure Legends for "FAK activity exacerbates disturbed flow-mediated atherosclerosis via VEGFR2-Cbl-NF-κB signaling"

Running title: Flow-mediated NF-κB signaling by FAK-VEGF2-Cbl axis

***Correspondence to:** Steve Lim, Ph.D., Department of Pathology, University of Alabama at Birmingham, 1825 University Blvd, Shelby 815, Birmingham, AL 35294

Key words: FAK, NF-κB, Cbl, VEGFR2, disturbed flow, Atherosclerosis
Word Count: 4745
Number of figures: 6

**Supplemental Information**

**Supplemental methods**

**Flow Experiment**

HUVECs were transfected with either HA-Cbl-WT or HA-Cbl-Y3F overnight. HUVECs were then subjected to oscillatory flow for either 0 or 30 min. Cells were lysed in SDS loading buffer and subjected to immunoblotting.

**Supplemental Figure legends**

**Supplemental Figure1.** HUVECs were starved for 12 h in 0.2% FBS DMEM with or without VEGF (5 ng/ml) prior to initiation of laminar flow (12 dynes/cm$^2$). Immunoblotting of active FAK (pY397), active NF-κB (pS536), active IKKα/β (pS176/177), IκBα, and either GAPDH or β-actin as loading control are shown.

**Supplemental Figure 2.** HUVECs were transfected with either HA-Cbl-WT or HA-Cbl-Y3F overnight. HUVECs were then subjected to oscillatory flow for either 0 or 30 min. Immunoblotting for VEGFR2, active pY397 FAK, HA Cbl, or GAPDH as loading control.

**Supplemental Figure 3.** HUVECs were starved for 12 h in 0.2% FBS DMEM with or without FAK-I (2.5 uM) prior to initiation of disturbed flow (5 dynes/cm$^2$). Immunostaining for EEA1 (green), VEGFR2 (red) and DNA (DAPI, blue).

**Supplemental Figure 4.** Partial carotid ligation (PCL) was performed on *Apoe-/-* mice and were treated with vehicle or FAK inhibitor (FAK-I, 30 mg/kg, twice daily) while on a western diet (WD) for 2 weeks. **(A)** Immunoblotting of lung lysates for FAK, active pY397 FAK, and either GAPDH or β-actin for loading controls. Immunostaining of carotid arteries for **(B)** pS536 NF-kB (Red) or **(C)** VCAM-1 (Red), or **(D)** CD68 (Green). **(B and C)** Endothelial cells were stained with vWF (Green). Nuclei with DAPI (Blue).

**Supplemental Figure 5.** Partial carotid ligation (PCL) was performed on *Apoe-/-;FAK-WT* and *Apoe-/-;FAK-KD* mice and fed a western diet (WD) for 2 weeks. Immunostaining of carotid arteries for **(A)** VCAM-1 (Red) or **(B)** pS536 NF-kB (Red). Endothelial cells stained with VWF (Green) and nuclei were stained with DAPI (blue).
