## Supplementary figures and images for "FAK activity exacerbates disturbed flow-mediated atherosclerosis via VEGFR2-Cbl-NF-κB signaling"

### Supplemental Figure 1

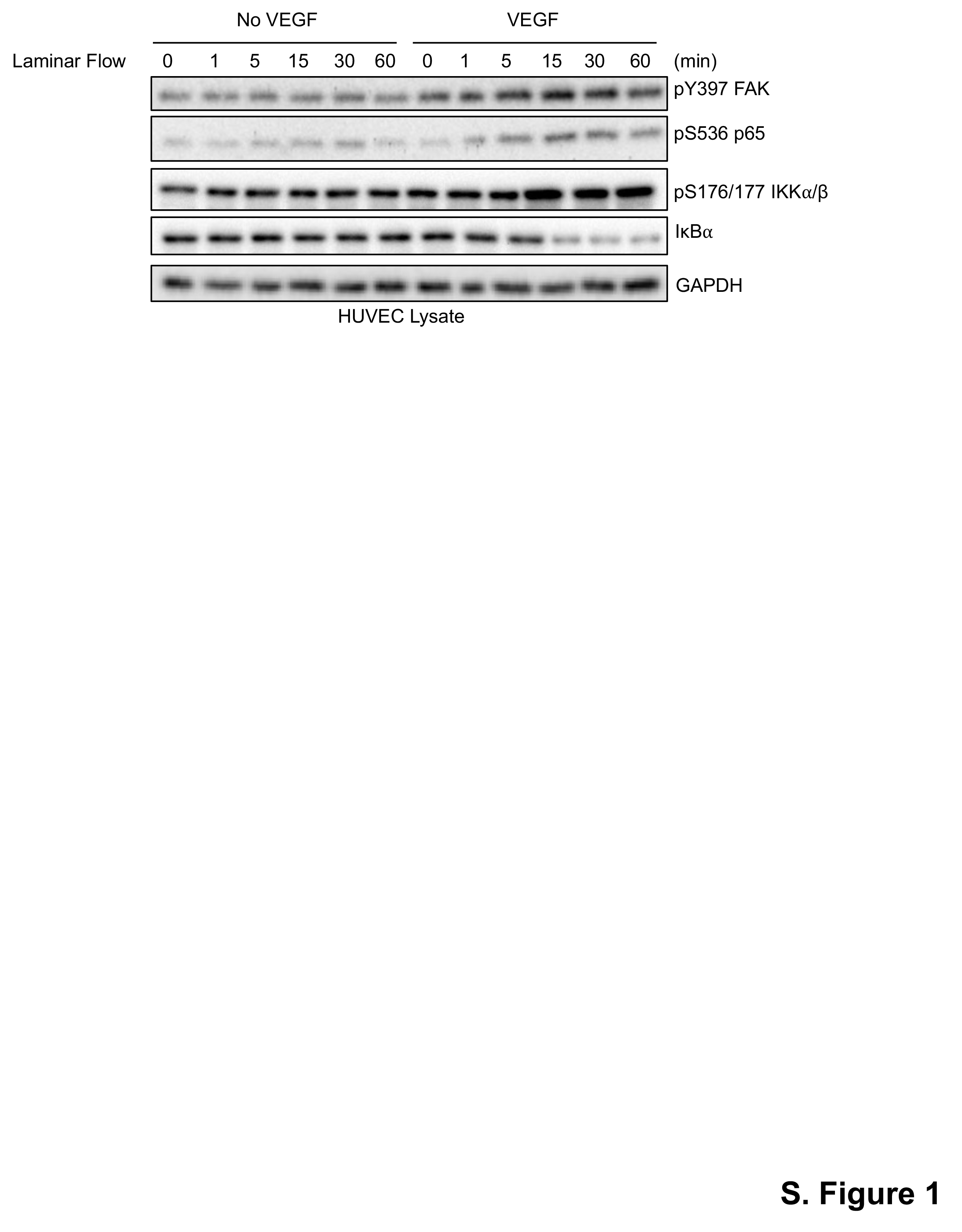

### Supplemental Figure 2

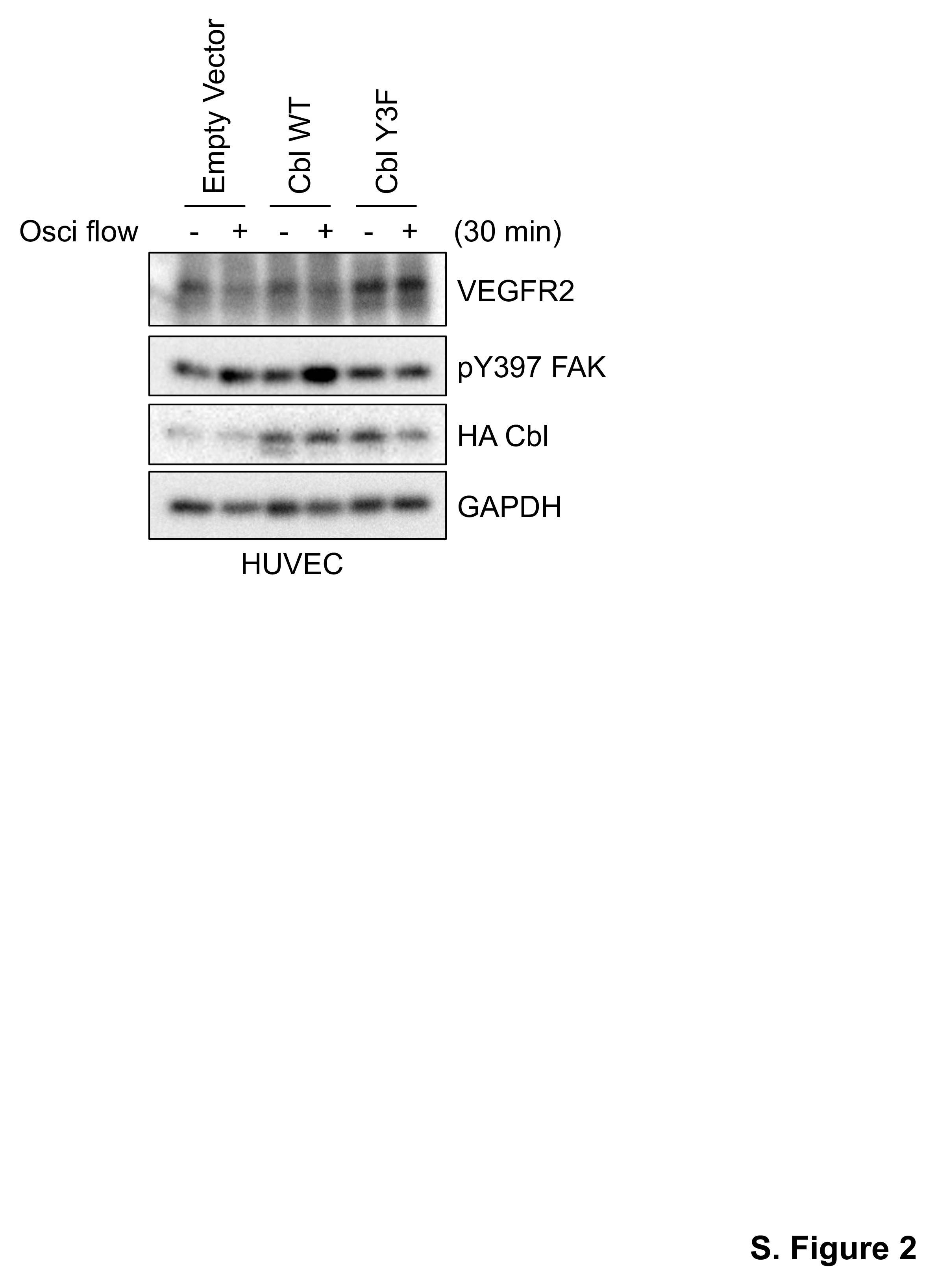

### Supplemental Figure 3

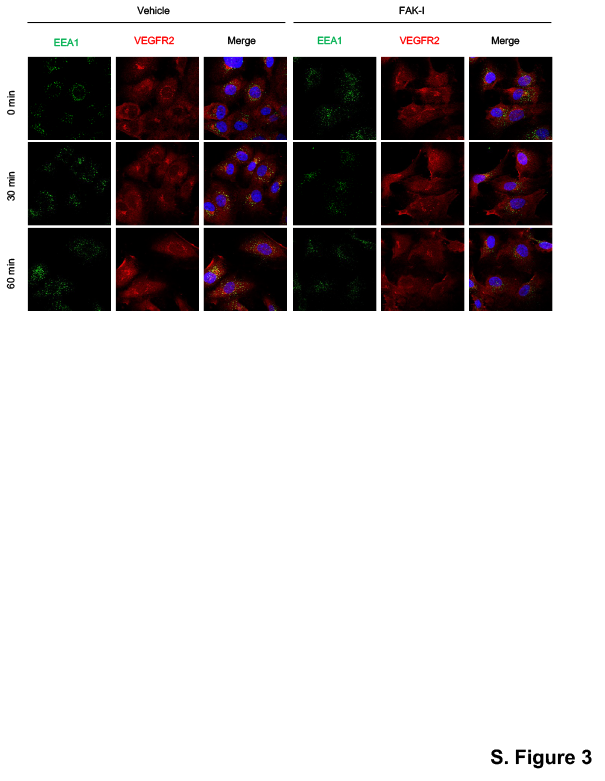

### Supplemental Figure 4

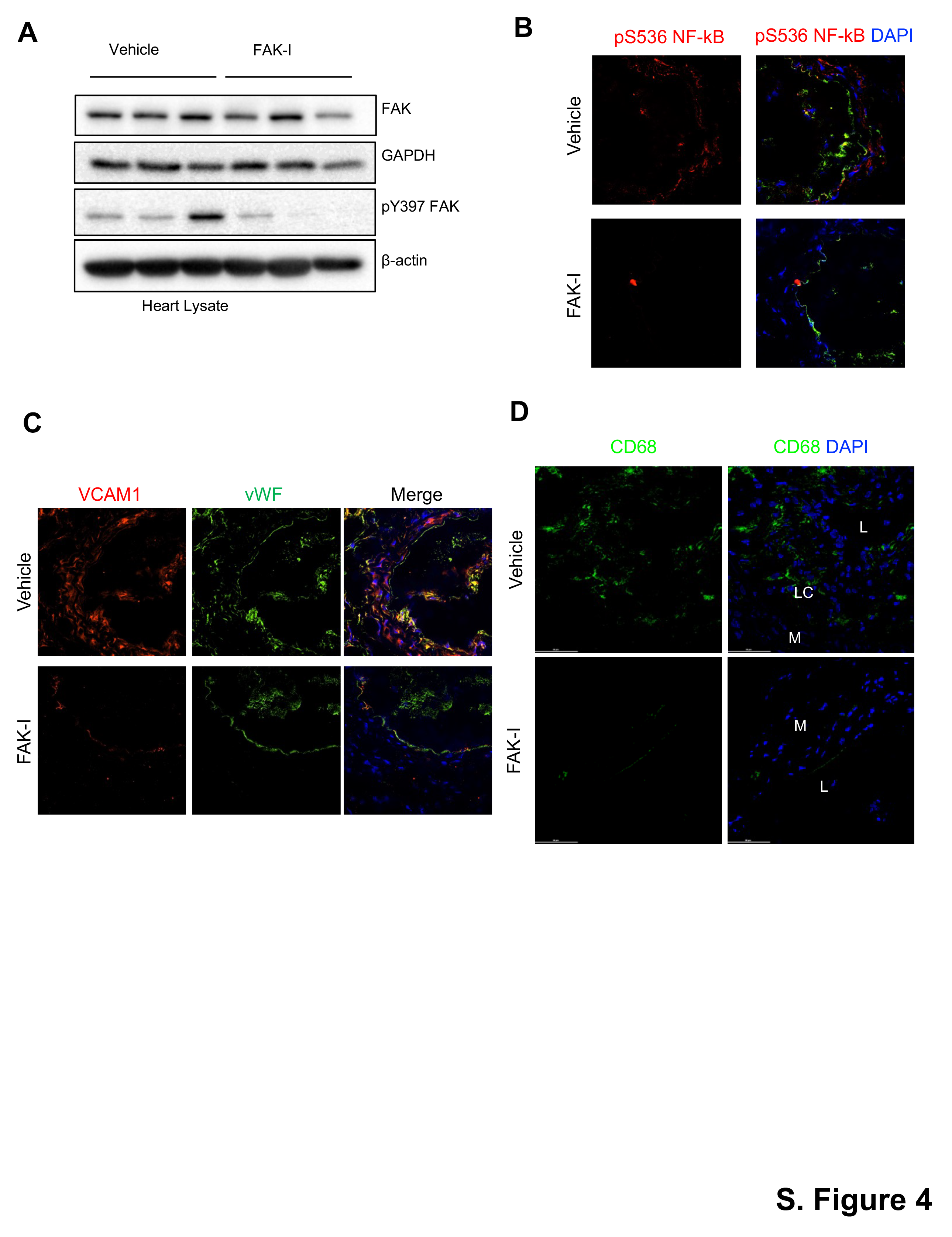

### Supplemental Figure 5

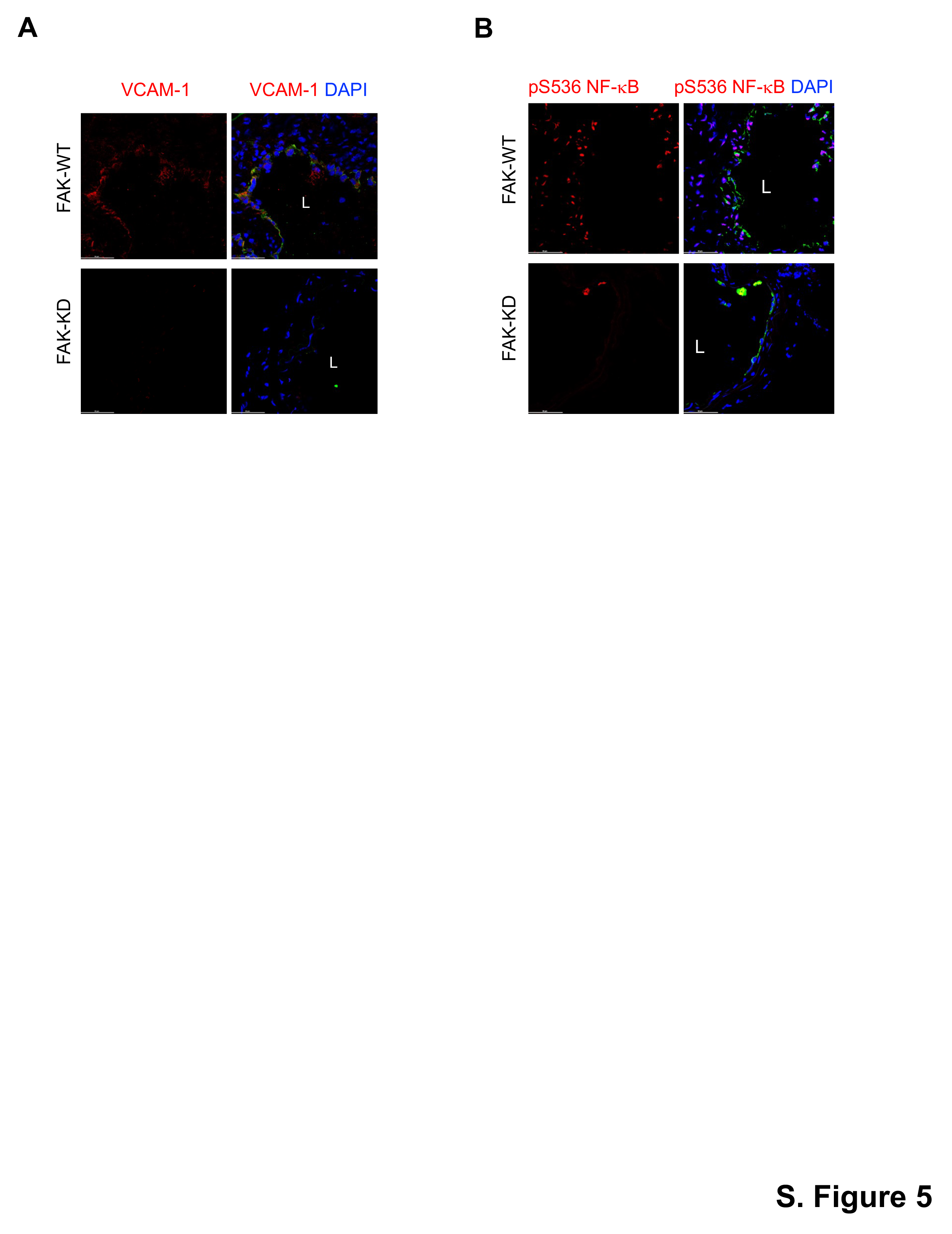
